# Correction of a pathogenic mutation in iPSCs derived from a patient with Christianson syndrome using CRISPR/Cas9 genome editing

**DOI:** 10.1101/2021.05.20.445025

**Authors:** Li Ma, Qing Wu, Michael Schmidt, Eric M. Morrow

## Abstract

*SLC9A6* (also termed *NHE6*) encodes the endosomal Na+/H+ exchanger 6 (NHE6). Pathogenic, loss-of-function mutations in *NHE6* cause the X-linked neurogenetic disorder Christianson syndrome (CS). We developed induced pluripotent stem cell (iPSC) lines derived from a patient with CS and from a biologically related control. The patient with CS contained the nonsense mutation c.1569G>A (p.(W523X)), which caused a significant reduction in *NHE6* mRNA and a lack of detectable NHE6 protein in CS iPSCs in comparison to control iPSCs. To establish a cell model for study of CS with an isogenic control, we corrected the c.1569G>A mutation to the *NHE6* reference genome sequence using CRISPR/Cas9-mediated homology directed repair knock-in methodology. Multiple subclonal lines were generated, and notably, NHE6 protein was expressed in all analyzed c.1569G>A (p.(W523X)) genome-corrected iPSC lines. This CS iPSC model together with the associated biologically related and isogenic control cell lines will serve as a valuable resource for both basic and translational studies in CS.

## 1. Introduction

Christianson syndrome (CS)^1^ is an X-linked neurogenetic disorder with clinical features including intellectual disability, postnatal microcephaly, ataxia, epilepsy, hyperkinesis, nonverbal status, and cerebellar atrophy (Christianson et al., 1999; Gilfillan et al., 2008; Morrow and Pescosolido, 2018; Padmanabha et al., 2017; Pescosolido et al., 2014; Schroer et al., 2010). CS is caused by pathogenic, loss-of-function mutations in *SLC9A6* (also termed *NHE6*), the gene encoding for the 12-transmembrane (TM)-domain, endosomal protein Na+/H+ exchanger 6 (NHE6) (Gilfillan et al., 2008; Morrow and Pescosolido, 2018).

Human induced pluripotent stem cells (iPSCs) have served as an important complement to animal models in translational studies targeting human genetic disease. IPSC resources have been widely used as a model for human diseases, including neurological and neuropsychiatric diseases such as Parkinson’s disease, Alzheimer’s disease, Huntington’s disease, schizophrenia, and autism (Brennand et al., 2011; Chae et al., 2012; Chen et al., 2020; Devine et al., 2011; Mungenast et al., 2016; Nehme and Barrett, 2020; Sullivan and Young-Pearse, 2017; Takahashi and Yamanaka, 2006; Tcw, 2019; Yagi et al., 2011).

In order to establish a patient-derived cellular model for the study of CS, we reprogrammed peripheral blood mononuclear cells from a male patient containing a c.1569G>A (p.(W523X)) mutation in *NHE6* and a clinical diagnosis of CS into iPSCs. We also generated important controls, including: 1) iPSCs from a biologically related male sibling who does not carry a pathogenic mutation in *NHE6*; and 2) isogenic iPSCs wherein the c.1569G>A mutation was corrected back to the reference sequence by using Clustered regularly interspaced short palindromic repeats (CRISPR)/CRISPR-associated protein 9 (Cas9)-mediated homology directed repair (HDR) knock-in methodology. Herein we report on an initial characterization of the CS iPSC lines containing a c.1569G>A (p.(W523X)) mutation in *NHE6* and the associated biologically related and isogenic control cell lines. These cell lines will serve as a valuable resource for both basic and translational studies in CS.

## 2. Materials and methods

### 2.1. Generation and maintenance of iPSCs

The Institutional Review Boards at Brown University and Lifespan Healthcare approved the human subjects research protocol. Informed consent was obtained from all participants or guardians of participants. The protocol for generation of iPSCs from peripheral blood mononuclear cells was adapted from Kunisato et al. (Kunisato et al., 2011); iPSC lines were classified based on the method used, either Sendai virus or lentivirus, for transducing vectors allowing for expression of reprogramming factors. The iPSC lines were cultured on Matrigel-coated plates (Corning #354277) under feeder-free conditions and maintained in mTeSR Plus media (StemCell Technologies #100-0276). Culturing medium was changed approximately daily. Enzyme-free passaging was performed using ReLeSR (StemCell Technologies #05872) according to the manufacturer’s instructions. Cell cultures were maintained at 37 °C in a humidified atmosphere of 95% air and 5% CO_2_. The iPSC lines generated as part of this study, including the genome-edited iPSC lines described below, will be deposited with an NIH repository for open sharing and distribution.

### 2.2. Genomic DNA extraction and Sanger sequencing

Genomic DNA from three control (line 404 subclones S1, S3, and S4) and four CS (line 403 subclones 1, S7, S8, and S9) iPSC lines was extracted using quick extract DNA extraction solution (Epicentre #QE09050), amplified by PCR, and sequenced using Sanger sequencing methods. Primers for PCR amplification of exon 12 of *NHE6* have been previously described (Pescosolido et al., 2014) and were as follows: forward 5’-AGAACGTGCTTTCTTGCTCTG -3’ and reverse 5’-TGGAAAGATATCCCTCAAAGTC -3’. Mutations were identified by verifying the chromatograms using Chromas Lite software and comparing the sequence data to the sequence reported for *NHE6* on the UCSC Genome Browser based on human genome build GRCh37/hg19 (http://genome.ucsc.edu) (Kent et al., 2002).

### 2.3. NanoString assay

Control (subclones 404-S1, -S3, and -S4) and CS (subclones 403-1, -S7, -S8, and -S9) iPSC lines were harvested and total RNA was collected using the RNeasy Mini Kit (Qiagen #74104). Messenger RNA (mRNA) amounts were analyzed using a custom-designed NanoString platform containing a probe that recognizes all three *NHE6* mRNA transcript variants (TVs), *NHE6* TV1 (NM_001042537), *NHE6* TV2 (NM_006359), and *NHE6* TV3 (NM_001177651). RNA counts were normalized to hybridization controls and to a set of housekeeping genes: *B2M, GAPDH, HPRT1, RPL13a*, and *POLR2A*. Subclones were analyzed individually as three technical replicates, with one replicate being excluded for a CS iPSC line due to poor quality. Combined data are presented for control iPSC lines and CS iPSC lines.

### 2.4. Western blotting and immunoprecipitation

For Western blotting, control, CS, and genome-corrected iPSCs were harvested and then lysed in lysis buffer (50 mM Tris-HCl, pH 7.8, 137 mM NaCl, 1 mM NaF, 1 mM NaVO_3_, 1% Triton X-100, 0.2% Sarkosyl, 1 mM dithiothreitol, and 10% glycerol) supplemented with protease inhibitor cocktail and phosphatase inhibitor for 30 min on ice. Cell lysates were separated by centrifugation at 13,200 rpm for 15 min at 4 °C, and the remaining supernatants were removed for further processing. Protein concentration was measured by BCA assay using the Pierce BCA Kit (Thermo Fisher #23225). For immunoprecipitation, aliquots of a custom-made rabbit anti-NHE6 antibody (C-terminal epitope: GDHELVIRGTRLVLPMDDSE, Covance #048) (Ouyang et al., 2013) were conjugated to Dynabeads Protein G (Thermo Fisher #10004D) at room temperature for 2 h according to the manufacturer’s instructions. Cell lysates were then incubated with anti-NHE6 antibody-conjugated beads overnight at 4 °C. The following day, the antibody-conjugated beads were gently pelleted, cell lysates were removed, and the antibody-conjugated beads were washed three times with phosphate-buffered saline with Tween 20 (PBST) wash buffer. The Dynabeads precipitates were then boiled in sample buffer at 95°C for 5 min before loading onto 4-12% SDS-PAGE gels (Novex #NP0321Box). Following separation of proteins by electrophoresis, gels were transferred to nitrocellulose membranes (Novex #LC2000). Western blots were performed using standard procedures (Lizarraga et al., 2021; Ouyang et al., 2019; Xu et al., 2017) and were analyzed with the Li-CoR Odyssey Imaging System. Proteins were detected using rabbit anti-NHE6 antibody (1:1,000 working dilution; Covance #048) (Ouyang et al., 2013) and mouse anti-α-Tubulin (1:5,000 working dilution; Sigma #T6074).

### 2.5. CRISPR/Cas9-mediated genome editing of CS iPSC line

CRISPR/Cas9-based technology was used to revert an iPSC line derived from a patient with CS harboring a c.1569G>A (p.(W523X)) mutation in exon 12 of the *NHE6* gene back to the reference sequence. Guide RNAs (gRNAs) were designed according to the CRISPOR web tool (http://crispor.tefor.net/) and were synthesized by Integrated DNA Technologies. Three gRNAs were tested and gRNA2 was finally selected as the best gRNA. A single-stranded oligodeoxyribonucleotide (ssODN) consisting of a 121-nucleotide ultramer (Integrated DNA Technologies) was used as template to repair the double-stranded break. The ssODN was designed to contain the desired reference *NHE6* sequence, flanked with homology arms to the targeted genomic region. The ssODN was also designed to contain three silent mutations, each in a protospacer adjacent motif (PAM) (PAM1, PAM2, and PAM3), to prevent re-targeting and recutting by Cas9. Introduction of a BtsIMutI restriction site (CAGTGTG) in introducing one of the silent mutations served as a means for identification of successfully targeted clones.

For gene editing, ∼4 × 10^5^ CS iPSCs containing a c.1569G>A (p.(W523X)) mutation were treated with Accutase (working dilution 1:3 in PBS; StemCell Technologies #07922,). Cells were dissociated to generate a suspension of single cells, which were resuspended in mTeSR Plus media with 10 µM Rho-associated protein kinase inhibitor (ROCKi, Tocris Bioscience #Y-27632) for 24 h. Cells were then pelleted, resuspended in ribonucleoprotein (RNP)/CRISPRMAX mix (Thermo Fisher #CMAX00003), and incubated for 5 min at room temperature before plating. For the RNP complex, 30 pmol TrueCut Cas9 Protein (Invitrogen #A36498) and 15 pmol gRNA were combined for a ratio of 2:1, and 50% of the RNP complex solution was mixed with a minimum of 40 pmol ssODN for transfection. After incubation for 72 h, genomic DNA was extracted from iPSCs, and PCR and Sanger sequencing were performed to determine the efficiency of knock in of the desired mutation (i.e., reversion of the c.1569G>A (p.(W523X)) mutation to the reference sequence).

To generate colonies derived from a single cell, the CS iPSCs transfected with the RNP/CRISPRMAX mix were diluted at ratios of 1:3,000 and 1:10,000, and plated on 10-cm dishes. Cells were grown on dishes, with medium of mTeSR Plus + ROCKi changed daily until colonies were visible. At this stage, the colonies were not necessarily derived from a single cell and were termed “pooled colonies.” Pooled colonies were isolated and transferred to 96-well Matrigel-coated plates (Corning #354277) for culturing and subsequent DNA extraction and Sanger sequencing. Pooled colonies for which sequencing chromatograms showed a high peak demonstrating correction of the c.1569G>A (p.(W523X)) mutation were then dissociated and sorted so as to have a single cell per well of a 96-well plate. Colonies derived from a single cell were then further cultured, and correction of the c.1569G>A (p.(W523X)) mutation was confirmed by Sanger sequencing.

### 2.6. Imaging of iPSCs

Cell colonies derived from iPSC lines in which the *NHE6* c.1569G>A (p.(W523X)) mutation had been corrected were imaged using a Nikon Eclipse TS100 microscope equipped with a Q Imaging QIClick 1.4-MP CCD monochrome microscope camera. Images were captured of colonies being visualized using a 4X objective.

### 2.7. Statistical analysis

Unpaired two-tailed Student’s *t* tests were performed for all group comparisons using GraphPad Prism 6 software. Data are presented as mean ± standard error of the mean (SEM). **** *p* < 0.0001.

## 3. Results

### 3.1. Validation of NHE6 sequence in CS and control iPSC lines

A family with one son affected with CS and carrying a c.1569G>A (p.(W523X)) mutation in *NHE6* and an unaffected, biologically related son was recruited for this study as a part of our larger CS study (Lizarraga et al., 2021; Pescosolido et al., 2014). The NHE6 protein has three domains: 1) an N-terminal domain; 2) a 12-TM domain, which includes the Na+/H+ exchanger domain; and 3) a cytoplasmic domain. The W523X mutation is located in TM domain 12 of the NHE6 protein (Krogh et al., 2001) (Fig. 1a). Upon Sanger DNA sequencing, control iPSC lines from the unaffected brother, all of which were reprogrammed using Sendai virus, showed a “G” at nucleotide position c.1569 (subclones 404-S1, -S3, and -S4, Fig. 1b, top panel). In contrast, in CS iPSC lines, the “G” was mutated to an “A” at nucleotide position c.1569 (Fig. 1b, middle and bottom panels). The c.1569G>A mutation was present in CS iPSC lines reprogrammed using either lentivirus (subclone 403-1, Fig. 1b, middle panel) or Sendai virus (subclones 403-S7, -S8, and -S9, Fig. 1b, bottom panel) (see also Supplementary Table 1). The c.1569G>A mutation is predicted to cause a p.W523X nonsense mutation in NHE6.

**Fig. 1.**
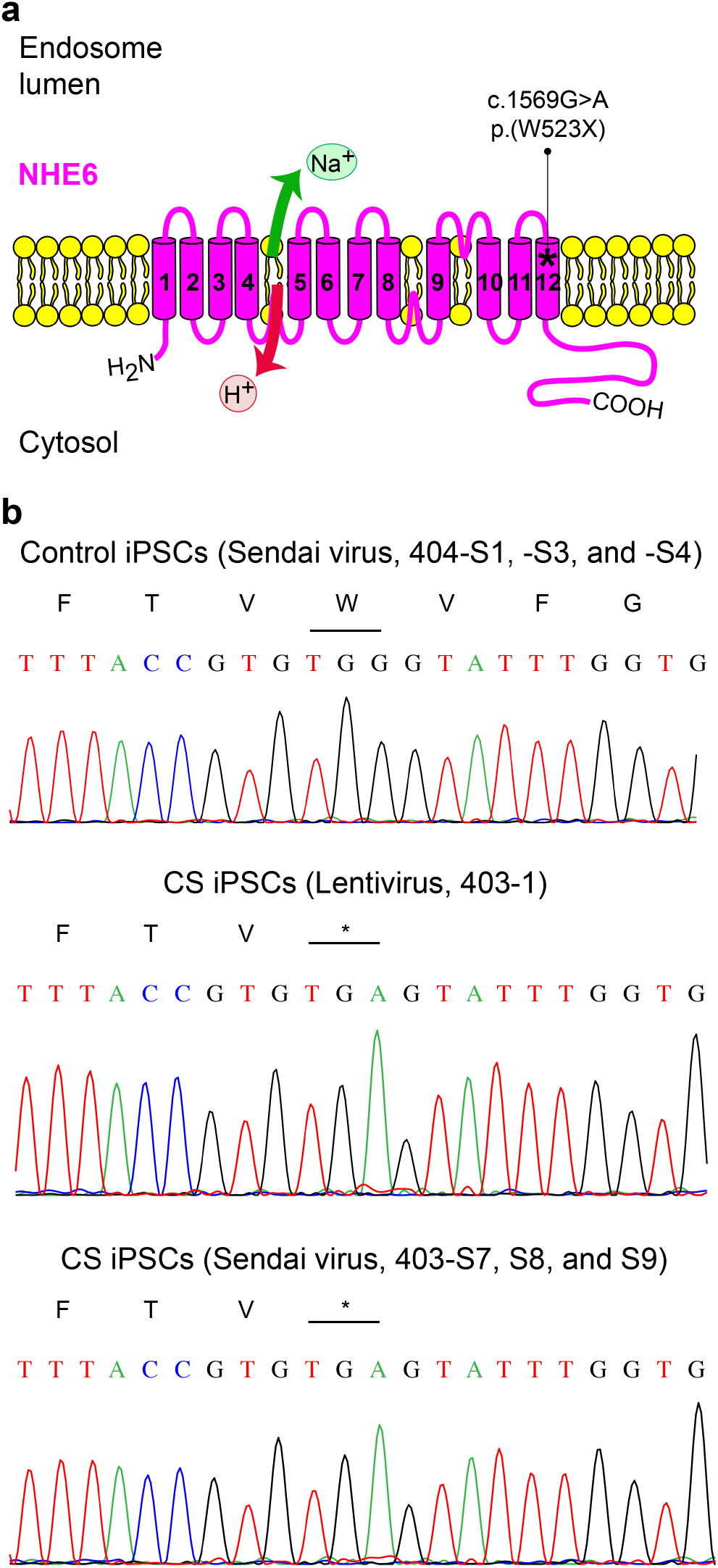
Sequence-based verification of an *NHE6* c.1569G>A (p.(W523X)) nonsense mutation in iPSCs derived from a patient with CS. **(a)** Schematic of the NHE6 protein. The approximate location of the c.1569G>A (p.(W523X)) mutation identified in the patient with CS from whom iPSCs were derived in the study described here is indicated. NHE6 TM domains were predicted using the transmembrane hidden Markov model (Krogh et al., 2001) for Ensembl transcript ENST00000370695 and protein ENSP00000359729 (Yates et al., 2016). **(b)** Sequencing chromatograms of the region covering the c.1569G>A (p.(W523X)) mutation for a genetically related, non-carrier brother control (top panel) and for the patient with CS (middle and bottom panels). IPSC lines were generated using either Sendai virus or lentivirus (as indicated) to transduce vectors allowing for expression of reprogramming factors. Genomic DNA was extracted from iPSCs, and exon 12 of *NHE6* was PCR-amplified and subsequently sequenced using Sanger sequencing methods. The location of the c.1569G>A (p.(W523X)) mutation is based on *NHE6* transcript NM_001042537.1 and NHE6 protein NP_001036002.1.

### 3.2. Absence of NHE6 mRNA and protein in NHE6 c.1569G>A (p.(W523X))-mutant iPSC lines

We first determined the effects of the c.1569G>A (p.(W523X)) mutation on NHE6 gene and protein expression. For gene expression, we used NanoString nCounter to determine the amounts of *NHE6* mRNA in samples (Control, subclones 404-S1, -S3, and -S4; CS, subclones 403-1, -S7, -S8, and -S9). Raw counts were normalized to hybridization controls and to a set of housekeeping genes, and normalized counts were compared between control and CS iPSC lines. *NHE6* mRNA is expressed as three different transcripts. We determined the amounts of *NHE6* mRNA in samples using a probe that recognizes all three TVs. As shown by the graph in Fig. 2a, the amount of *NHE6* mRNA was significantly reduced in CS iPSCs as compared to control iPSCs (*p* < 0.0001).

**Fig. 2.**
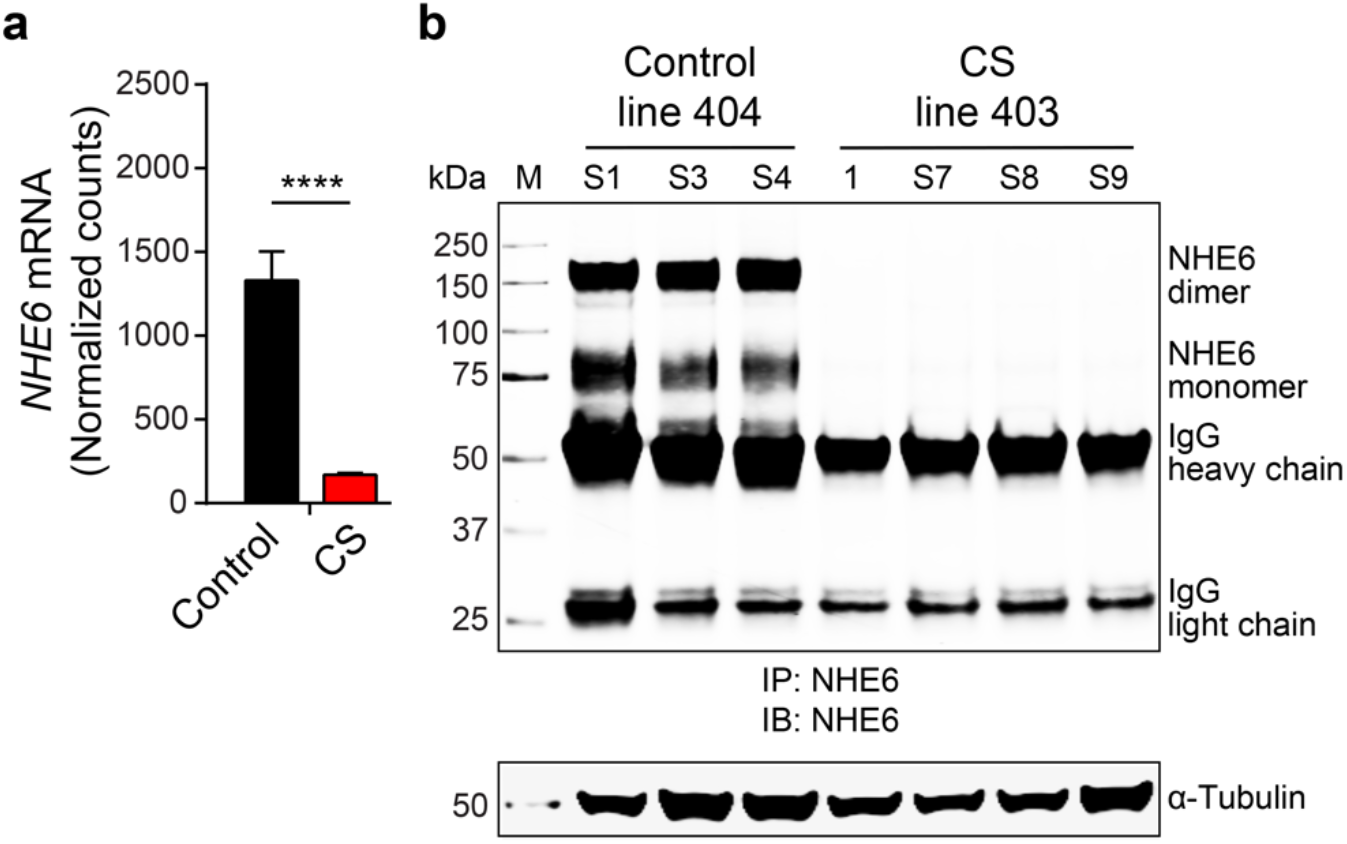
Analysis of NHE6 mRNA and protein expression in CS and paired control iPSC lines. **(a)** Graph depicting quantification of *NHE6* mRNA expression in CS (red bar) and paired control (black bar) iPSC lines based on NanoString data. *NHE6* mRNA expression was analyzed using a probe that recognizes all three *NHE6* TVs. Expression of *NHE6* mRNA was normalized to hybridization controls and to a set of housekeeping genes: *B2M, GAPDH, HPRT1, RPL13a*, and *POLR2A*. Control (subclones 404-S1, -S3, and -S4) and CS (subclones 403-1, -S7, -S8, and -S9) iPSC lines were analyzed individually as three technical replicates for each subclone, with one replicate being excluded for a CS iPSC line due to poor quality. Control: *n* = 9; CS: *n* = 11. **(b)** Western blot of lysates from CS and paired control iPSC lines, as labeled. Lysates were immunoprecipitated using our custom-made anti-NHE6 antibody and subsequently analyzed by Western blot using the same antibody. Blotting against α-Tubulin was used as a loading control. Data are presented as mean ± SEM. Unpaired two-tailed Student’s *t* test was used. ^****^ *p* < 0.0001.

The effect of the c.1569G>A (p.(W523X)) mutation on NHE6 protein expression was determined by Western blot using our custom-made anti-NHE6 antibody (Fig. 2b). NHE6 protein could be detected in control iPSCs as both a dimer and monomer at the predicted molecular weights (Fig. 2b, left three lanes). In contrast, in CS iPSCs, NHE6 protein was not detected, regardless of method of reprogramming (Fig. 2b, right four lanes). Taken together, the data support a significant reduction in *NHE6* mRNA in CS iPSCs in comparison to control iPSCs and a lack of detectable amounts of NHE6 protein in CS iPSCs.

### 3.3. Correction of NHE6 c.1569G>A (p.(W523X)) mutation using CRISPR/Cas9-mediated genome editing

The control iPSC lines for which results are reported above were derived from the unaffected brother of the patient with CS. Due to genetic variability between siblings, the control and CS iPSC lines likely differ at sites in addition to the c.1569G>A (p.(W523X)) mutation. To generate iPSC lines isogenic to a CS iPSC line, CRISPR/Cas9-mediated HDR knock-in methodology was used to correct the c.1569G>A (p.(W523X)) mutation back to the *NHE6* reference sequence. The c.1569G>A (p.(W523X)) mutation is located in exon 12 of *NHE6* (Fig. 3a). CS iPSC line 403-S7 was randomly selected as template for genome editing. Three gRNAs were designed to target exon 12 of *NHE6* (Fig. 3b), and an ssODN with the targeted A>G mutation at the exact center was designed for HDR (Fig. 3c, red text). To prevent re-targeting and recutting by Cas9, the ssODN was also designed to contain three silent mutations, each in a PAM (PAM1, PAM2, and PAM3) (Fig. 3c, blue text). A BtsIMutI restriction site (CAGTGTG) was introduced in PAM2 and served as a means for identification of successfully targeted clones. After testing, gRNA2 was selected as the best gRNA. Following transfection of the RNP/Cas9 complex together with ssODN and subsequent colony selection, Sanger sequencing was used to confirm correction of the c.1569G>A (p.(W523X)) mutation back to the *NHE6* reference sequence (Fig. 3d and Supplementary Table 1) (see Materials and methods for details). From 24 sequenced polyclonal colonies, two clones were identified showing the A>G correction, one of which was expanded so as to generate clones derived from a single cell. Overall, 14 clonal genome-corrected CS iPSC lines derived from a single cell were generated. Three polyclonal genome-corrected CS iPSC lines derived from two or more clones, but showing homogeneity for the A>G correction by sequencing, were also generated (Supplementary Table 1).

**Fig. 3.**
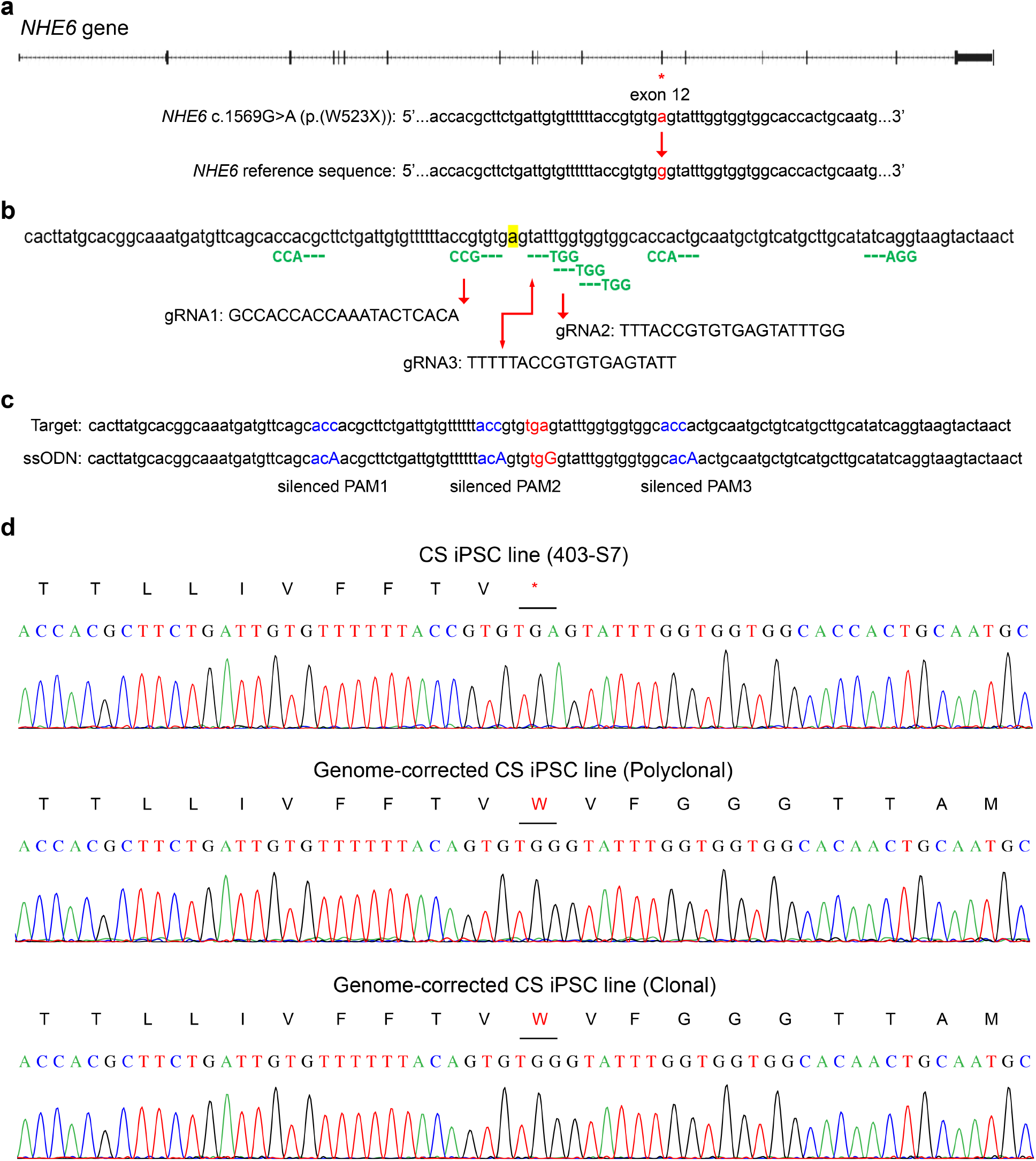
Correction of the c.1569G>A (p.(W523X)) mutation in CS iPSCs to the *NHE6* reference sequence using CRISPR/Cas9-mediated genome editing. **(a)** Schematic of the *NHE6* gene [adapted from UCSC Genome Browser, GRCh38/hg38, http://genome.ucsc.edu (Kent et al., 2002)] and immediate sequence surrounding the c.1569G>A (p.(W523X)) mutation, located in exon 12. The red asterisk indicates the location of exon 12. The aim of the genome editing was to correct the “a” nucleotide mutation back to a “g” nucleotide, as in the *NHE6* reference sequence. **(b)** Depiction of mutant *NHE6* target sequence (mutation site highlighted in yellow), PAMs of seven candidate gRNAs (green text), and three selected gRNAs. The CRISPOR web tool was used to design gRNAs for genome editing. Three dotted lines on the left of a PAM indicate the gRNA was designed in the same direction as the sense strand; three dotted lines on the right of a PAM indicate the gRNA was designed in the same direction as the antisense strand. The three indicated gRNAs were selected after analyzing the gRNA “specificity score” and “off target” information. After testing, gRNA2 was selected as the best gRNA. **(c)** Depiction of the target sequence (top) and ssODN (bottom). The mutant and corrected codons are labeled in red text, with the corrected “G” nucleotide capitalized in the ssODN. Silent mutations were introduced into three PAMs (blue text) to prevent recutting by Cas9. The “acc” codons were mutated to “aca”; the mutated “A” nucleotide is capitalized in the ssODN. **(d)** Sequencing chromatograms of DNA extracted from the CS iPSC line used as template for genome editing (subclone 403-S7, top panel) and from a polyclonal colony (middle panel) and a clonal colony (bottom panel) in which the c.1569G>A (p.(W523X)) mutation had been corrected to the *NHE6* reference sequence. The stop codon in the CS iPSC line is indicated with an asterisk.

### 3.4. Recovery of NHE6 protein expression in genome-corrected CS iPSCs and display of typical stem cell morphology

After sequencing to confirm correction of the c.1569G>A (p.(W523X)) mutation at the DNA level, Western blotting was performed to confirm recovery of NHE6 protein expression (Fig. 4a). Cell lysates were extracted from harvested iPSCs from an unaffected sibling control iPSC line, the CS iPSC line used for genome editing, and 14 genome-corrected CS iPSC lines (12 clonal and 2 polyclonal). Using our custom-made anti-NHE6 antibody, NHE6 protein (monomeric and dimeric forms) was detected in lysates from the control iPSC line and from the genome-corrected CS iPSC lines, regardless of clonal or polyclonal line, but not in lysate from the CS iPSC line used for genome editing (Fig. 4a).

**Fig. 4.**
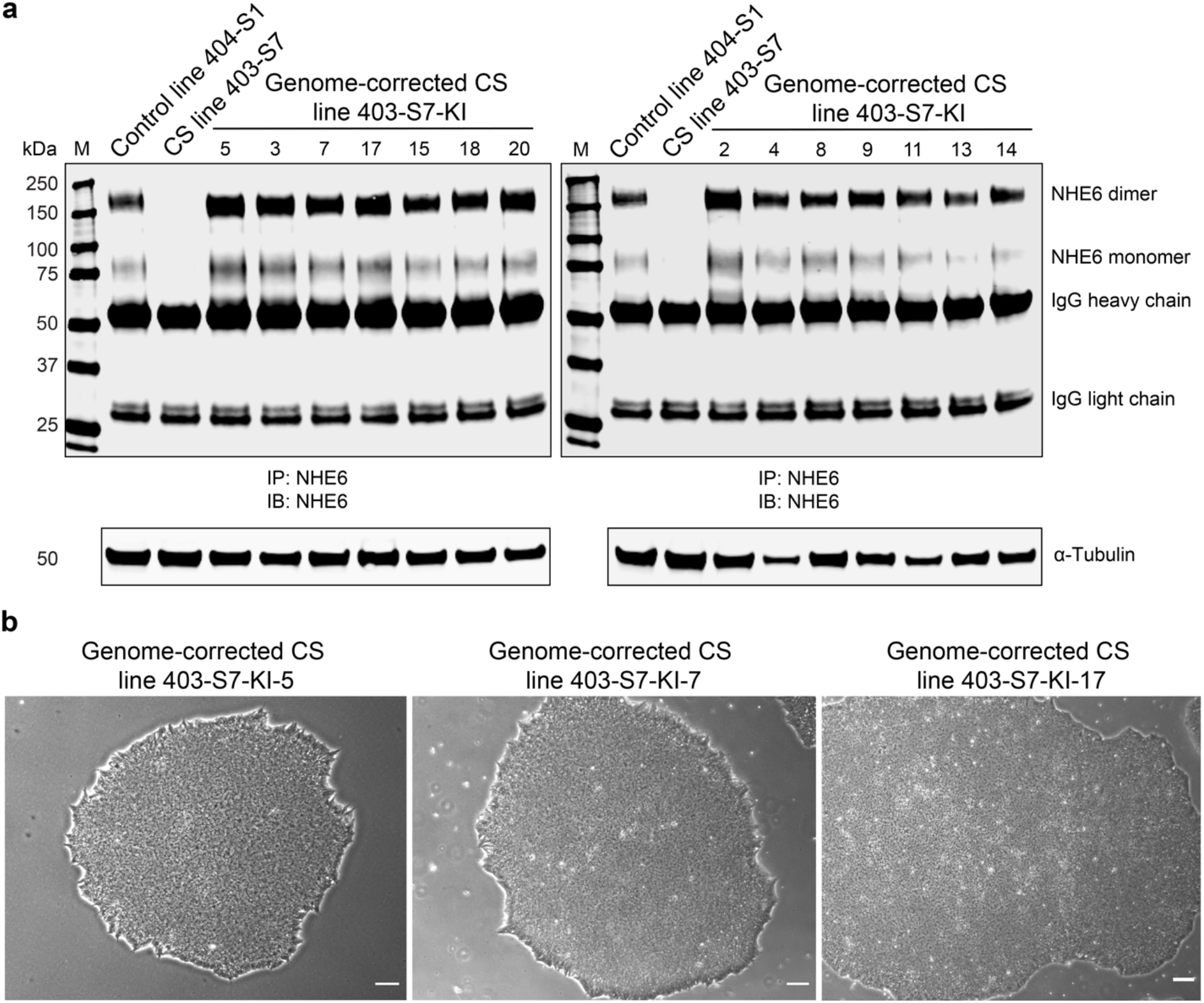
Expression of NHE6 protein in genome-corrected CS iPSCs and imaging of clonal cell colonies. **(a)** Western blot of lysates from paired control, CS, and genome-corrected CS iPSC lines, as labeled. Lines 15 and 18 are reflective of polyclonal colonies, whereas all other lines are reflective of clonal colonies. All lines showed correction of A>G. Lysates were immunoprecipitated using our custom-made anti-NHE6 antibody and subsequently analyzed by Western blot using the same antibody. Blotting against α-Tubulin was used as a loading control. **(b)** Brightfield microscopy images of three representative clonal colonies of genome-corrected CS iPSCs derived from a single cell. Scale bar, 100 µM.

The morphology of genome-corrected CS iPSCs from the different clonal and polyclonal lines was observed under brightfield microscopy. Images of selected representative examples of clonal colonies are shown in Fig. 4b. In general, the genome-corrected CS iPSCs displayed classic pluripotent stem cell morphology with a high nucleus to cytoplasm ratio and grew as high-density monolayers of tightly packed cells.

## 4. Discussion

In this study, we report on iPSC lines derived from a patient with CS containing an *NHE6* c.1569G>A (p.(W523X)) mutation, as well as on two important sets of control lines: 1) iPSC lines derived from a biologically related male sibling who does not carry a pathogenic mutation in *NHE6*; and 2) isogenic iPSC lines wherein the c.1569G>A mutation was corrected back to the reference *NHE6* sequence by using CRISPR/Cas9-mediated HDR knock-in methodology. Using NanoString nCounter, we found a significant reduction in *NHE6* mRNA in CS iPSCs in comparison to control iPSCs. Consistently, results from Western blot analysis revealed a lack of detectable amounts of NHE6 protein in CS iPSCs, regardless of reprogramming method (lentivirus or Sendai virus). Notably, in genome-corrected CS iPSC lines, which were confirmed by Sanger sequencing, expression of NHE6 protein was recovered. Cells from these lines also displayed classic stem cell morphology and colony growth characteristics.

The c.1569G>A (p.(W523X)) mutation is located in exon 12 of the *NHE6* gene and in TM domain 12 of the NHE6 protein. The reduction in *NHE6* mRNA and lack of NHE6 protein in CS iPSCs containing this mutation support that it is a loss-of-function mutation. Although not explored in detail here, based on our experimental results from other CS iPSC lines with nonsense mutations (Lizarraga et al., 2021), we anticipate that the c.1569G>A (p.(W523X)) mutation is subject to nonsense-mediated mRNA decay.

For studies based in use of iPSCs, isogenic controls are ideal as a means to reduce the confounds of genomic variation present when using unrelated controls and even in biologically related controls. CRISPR/Cas9-mediated genome editing is a powerful tool not only for inducing mutations but also for making corrections. Here we used CRISPR/Cas9-mediated HDR knock-in methodology to correct the c.1569G>A (p.(W523X)) mutation back to the *NHE6* reference sequence. The CRISPOR web tool (http://crispor.tefor.net/) was used to design gRNAs. Some difficulties were encountered in choosing an appropriate PAM due to the potential for recutting by Cas9 and an inability to introduce functional silent mutations. For genome editing, we transfected the RNP complex (Cas9 and gRNA) together with the ssODN into CS iPSCs. RNP complexes at Cas9:gRNA ratios of 5:1 and 2:1 were tested, and it was determined that a 2:1 ratio was more optimal. In optimizing conditions, we also tested cotransfection of different amounts of RNP complex (100%, 75%, 50%, or 25%) and ssODN; here, 50% of RNP complex was most optimal for the cotransfection. Finally, rather than the supplier-recommended forward transfection protocol, we used a reverse transfection protocol. The CRISPR/Cas9-mediated genome editing approach for correcting the c.1569G>A (p.(W523X)) mutation was successful; however, the HDR efficiency was somewhat low, with only 2 of 24 selected polyclonal colonies showing the A>G correction upon Sanger sequencing.

In summary, we have generated iPSC lines derived from a patient with CS containing an *NHE6* c.1569G>A (p.(W523X)) mutation and from a biologically related male sibling who does not carry a pathogenic mutation in *NHE6*. Further, using CRISPR/Cas9-mediated HDR knock-in methodology, we generated isogenic iPSC lines wherein the c.1569G>A mutation was corrected back to the reference *NHE6* sequence. These lines have been validated at the levels of DNA sequence (Sanger sequencing), mRNA (NanoString nCounter), and protein (Western blot). These CS and control iPSC lines will serve as a useful resource and *in vitro* cell model for the study of molecular and cellular mechanisms of CS disease pathology and for treatment development. As such, we will deposit the lines with an NIH repository for open sharing and distribution.

## Author contributions

Li Ma: Formal analysis, Investigation, Writing – Original Draft, Visualization. Qing Wu: Formal analysis, Writing – Review & Editing. Michael Schmidt: Investigation, Data curation. Eric M. Morrow: Conceptualization, Methodology, Formal analysis, Writing – Review & Editing, Supervision, Funding acquisition.

## Acknowledgements

We thank the family who participated in this study. The study was supported by NIMH R01MH105442, NIMH R21MH115392, and NINDS/NIA R01NS113141 and by the Hassenfeld Child Health Innovation Institute at Brown University (E.M.M.). We thank the Brown University/Rhode Island Hospital Flow Cytometry and Sorting Facility and the Cincinnati Children’s Hospital Medical Center Pluripotent Stem Cell Facility for their services. We acknowledge Heather M. Thompson, Ph.D. of Brown University for her support in manuscript preparation and Li Li, Ph.D. of the Harvard Stem Cell Institute for technical advice and discussions.

## Supplementary data

**Supplementary Table 1.** Control, CS, and genome-corrected CS iPSC lines.

| Line | Subclone IDs | Reprogramming Method |  |
| --- | --- | --- | --- |
| Control | 404-S1 | Sendai virus transduction method |  |
|  | 404-S3 | Sendai virus transduction method |  |
|  | 404-S4 | Sendai virus transduction method |  |
| CS (c.1569G>A [p.(W523X)]) | 403-1 | Lentivirus transduction method |  |
|  | 403-S7 | Sendai virus transduction method | *Template for genome editing |
|  | 403-S8 | Sendai virus transduction method |  |
|  | 403-S9 | Sendai virus transduction method |  |

| Line | Subclone IDs | Clonal Status |
| --- | --- | --- |
| Genome-corrected CS | 403-S7-KI-1 | Clonal clone |
|  | 403-S7-KI-2 | Clonal clone |
|  | 403-S7-KI-3 | Clonal clone |
|  | 403-S7-KI-4 | Clonal clone |
|  | 403-S7-KI-5 | Clonal clone |
|  | 403-S7-KI-6 | Clonal clone |
|  | 403-S7-KI-7 | Clonal clone |
|  | 403-S7-KI-8 | Clonal clone |
|  | 403-S7-KI-9 | Clonal clone |
|  | 403-S7-KI-10 | Clonal clone (very small portion of A peak) |
|  | 403-S7-KI-11 | Clonal clone |
|  | 403-S7-KI-12 | Clonal clone (very small portion of A peak) |
|  | 403-S7-KI-13 | Clonal clone |
|  | 403-S7-KI-14 | Clonal clone |
|  | 403-S7-KI-15 | Possible 2 or more clones |
|  | 403-S7-KI-16 | Clonal clone (very small portion of A peak) |
|  | 403-S7-KI-17 | Clonal clone |
|  | 403-S7-KI-18 | Possible 2 or more clones |
|  | 403-S7-KI-19 | Possible 2 or more clones |
|  | 403-S7-KI-20 | Clonal clone |

## Footnotes

1 Cas9: CRISPR-associated protein 9; CRISPR: Clustered regularly interspaced short palindromic repeats; CS: Christianson syndrome; gRNA: Guide RNA; HDR: Homology directed repair; iPSC: Induced pluripotent stem cell; mRNA: Messenger RNA; NHE6: Na+/H+ exchanger 6; PAM: Protospacer adjacent motif; PBS: Phosphate-buffered saline; PBST: Phosphate-buffered saline with Tween 20; RNP: Ribonucleoprotein; ROCKi: Rho-associated protein kinase inhibitor; SDS-PAGE: Sodium dodecyl sulfate-polyacrylamide gel electrophoresis; ssODN: Single-stranded oligodeoxyribonucleotide; TM: Transmembrane; TV: Transcript variant.

